# Glial cell mechanosensitivity is reversed by adhesion cues

**DOI:** 10.1101/865303

**Authors:** C. Tomba, C. Migdal, D. Fuard, C. Villard, A. Nicolas

**Affiliations:** University of Geneva; Univ. Grenoble Alps, CNRS, LTM, 38000 Grenoble, France & Univ. Grenoble Alps, CEA, CNRS, Inserm, BIG-BCI, 38000 Grenoble, France & Univ. Grenoble Alps, CEA, Inserm, BIG-BGE, 38000 Grenoble, France; Univ. Grenoble Alps, CNRS, LTM, 38000 Grenoble, France; CNRS - Institut Néel; Univ. Grenoble Alps CNRS

## Abstract

Brain tissues demonstrate heterogeneous mechanical properties, which evolve with aging and pathologies. The observation in these tissues of smooth to sharp rigidity gradients raises the question of brain cells responses to both different values of rigidity and their spatial variations. Here, we use recent techniques of hydrogel photopolymerization to achieve stiffness structuration down to micrometer resolution. We investigate primary neuron adhesion and orientation as well as glial cell adhesive and proliferative properties on multi-rigidity polyacrylamide hydrogels presenting a uniform density of adhesive molecules. We first observed that neurons grow following rigidity gradients. Then, our main observation is that glial cell adhesion and proliferation can be enhanced on stiff or on soft regions depending on the adhesive coating of the hydrogel, i. e. fibronectin or poly-L-lysine/laminin. This behavior was unchanged in the presence or not of neuronal cells. In addition, and contrarily to other cell types, glial cells were not confined by sharp, micron-scaled gradients of rigidity. Our observations suggest that their mechanosensitivity could involve adheison-related mechanosensitive pathways that are specific to brain tissues.

**SIGNIFICANCE:** By growing primary brain cells on 2D multi-rigidity polyacrylamide hydrogels, we show that favorable culture conditions for glial cells switch from stiff to soft substrates when changing the adhesive ligands from fibronectin to poly-L-lysine/laminin. Together with neurons, glial cells thus provide a unique example where soft is preferred to stiff, but unlike neurons, this preference can be reversed by changing the nature of the coating. We additionally show that contrarily to other cell types, glial cells are deformed by subcellular gradients of rigidity but cannot be confined by these rigidity gradients. These observations point that glial cell use a very specific, integrin-related machinery for rigidity sensing.

## INTRODUCTION

Neurons, macroglial cells (astrocytes and olygodendrocytes) and microglial cells constitute the three major categories of brain cells. Among glial cells, astrocytes, whose relative number increases with brain complexity (1), are the most abundant (2). Since astrocytes have been recognized for their active and specific functions in brain computation (3), their multiple roles beyond providing cohesion of the brain tissue (glia coming from the greek for “glue”) is increasingly understood (4). In addtion, together with the other types of glial cells, astrocytes are involved in the response to traumatic brain insults (5, 6) and in cancers (7) from their intrinsic ability to proliferate and secrete tumorigenic factors. It is therefore crucial to better understand their response to the different stimuli (chemical, topographical, mechanical) provided by their microenvironments. Indeed, at the cell level, these cues trigger various cellular adaptations concerning adhesion, polarization, migration or proliferation among others (8, 9). For the last two decades, the study of cell response to the mechanical environment has attracted increasing attention (10), both in vivo (11, 12) and in vitro (13–15). Mechanical cues were for example shown to drive the epithelial to mesenchymal transition in vitro (16), and a recent in vivo study also demonstrated that tumor propagation can be mediated by mechanical signaling pathways (17). Links between the mechanical properties of the brain tissues and brain functions were also evidenced in vivo (18).

Significant changes on the cell structure and functions in response to the rigidity of the extracellular matrix have been observed in many cell types like fibroblasts or HeLa cells, or even stem cells including neural stem cells (19). Neurons have also shown sensitivity to rigidity. Neurites have been reported to grow faster on matrices below 1 kPa compared to stiffer substrates, either in 2D and in 3D (20–27). Consequently, the mechanosensitivity of the growth cone has been investigated (22, 24, 27–30), and neuron rigidity sensing was shown to be integrin-dependent (24). Glial cells however have received less attention. They were also shown to be sensitive to rigidity, but unlike neurons, they display better adhesion and larger spreading area on stiff substrates (31). They were also shown to display higher proliferation rates on the stiff substrates rather than on the soft ones (5, 21, 31).

Aside being among the softest tissues (32), brain tissues demonstrate heterogeneous elastic properties either in between the different tissues that compose the brain, with Young’s modulus ranging from few hundreds of Pa in the white matter (33–35) to several tens of kPa when addressing myelinated fibres (36) and pituitary gland tissues (37), as well as inside individual anatomical regions (37, 38). Smooth, millimeter-sized (39) and sharp, micron-sized (37, 40, 41) gradients of rigidity have been reported in brain tissues. Additionally, the stiffness of the brain tissues undergoes changes during brain development (42) and in pathological conditions (38, 40). The brain stiffness values and the presence of rigidity gradients at the scale of a few micrometers raises fundamental issues about brain cell mechanosensitivity. They in addition provide an instrumental challenge for in vitro studies.

In the present work, we address the contribution of the stiffness of the extra-cellular matrix (ECM) on the adhesion, growth and proliferation of primary neurons and glial cells (i.e. mostly astrocytes) dissociated from mouse embryo cortex and hippocampal tissues. In this aim, we cultured them on multi-rigidities hydrogels with apposed areas characterized by two distinct values of rigidity (1.1 and 42 kPa for most designs, but also 29 and 75 kPa) which replicate the range of rigidities found in the brain tissue. Moreover, the size of these areas could be tuned from the centimeter down to the micrometer scale using a recent technique of hydrogel stiffness structuration. In this way, we were able to address primary neuron and astrocytes response to sharp gradients of rigidity, and to directly compare cell responses to pairs of rigidity in identical culture conditions and in the presence of the soluble cues secreted by the cells on each rigidity condition. We then addressed primary neuron and glial cell response to multi-rigidity composite matrices presenting sharp gradients of rigidity, thus mimicking the in vivo situation where a population of cells from identical type is facing different rigidities because of the architecture of the ECM. Neuron and glial cells response, in terms of adhesion and proliferation, was systematically quantified in dependence on the nature of the adhesive ligands in the ECM, the composition of the culture medium, and the initial composition in cells in the culture (pure glial cells versus mixed neuron-glial cells co-cultures). We first observed that neuronal cells respond to sharp gradients of rigidity similarly to what was reported on smooth ones (43), with neurites growing parallel to the gradient of rigidity. Then, our main finding is that glial cells showed a differential response to rigidity depending whether adhesive ligands were fibronectin or poly-L-lysine/laminin: adhesion and proliferation were enhanced on the stiff substrates when cultured on fibronectin, while soft substrates were preferred on poly-L-lysine/laminin coating. These observations, that were not dependent on mixed versus pure culture conditions, are a unique demonstration that some integrin subtypes may favor cell adhesion or proliferation on soft matrices.

## MATERIALS AND METHODS

### Substrate fabrication, functionalization and characterization

Multi-rigidities polyacrylamide gels were fabricated from the protocols of photopolymerization. The monomer solution for polyacrylamide is prepared with a UV sensitive initiator. 2 mg of Irgacure 819 (Ciba Specialty Chemicals Inc., Basel, Switzerland) is dissolved into 10 µl of propylamine (240958, Sigma-Aldrich, St. Louis, USA) at 52°C for 10 min. 490 µl of deionized water, 250 µl of a 40% solution of acrylamide and 250 µl of a 2% solution of N, N-methylene-bis-acrylamide (Bio-Rad, Herculers, USA) are added, leading to a 10%-0.5% mixture of monomers.

Grey level masks are engineered using optical photolithography and etching on a microscope glass slide. The glass is first washed into a Piranha solution (H2O2: H2SO4) with concentration 1:2 for 10 min. Then, 1 nm of titanium and the desired thickness of chromium are deposited onto it with a Plassys type electron gun. The thickness of the chromium layer determines the transmission coefficient of the grey level (Fig. S1), and consequently the reticulation rate of the hydrogel (44). AZ1512HS resist (Clariant, Muttenz, Switzerland) is spun onto the metal deposit at 3000 rpm for 30 s to reach a 600 nm thickness. It is then illuminated through a black and white master lithographic mask that reproduces the patterns that are to be transferred in the hydrogel. The resist pattern is developed with 1:1 AZ-developer:de-ionized water mixture for 1 min. Etching is performed in a DPS type etching reactor, using a chlore:oxygen (2:1) plasma. The resist is removed by exposing it 30 s to an oxygen plasma in the DPS reactor. The grey level mask is then rendered hydrophobic by grafting a fluorinated silane onto it: the grey level mask is immersed into a 1‰ solution of Optool (DSX, Daikin, Pierre-Bénite, France) in perfluorohexane for 1 min. It is allowed to react 1 h in water vapor at 65°C. It is then rinsed for 10 min in perfluorohexane. The grey level chromium mask is then fitted with 40 µm thick wedges on its edges.

20 mm diameter coverslips are used as a support to the hydrogel. They are first cleaned in a 0.1 M NaOH solution for 10 min, then rinsed in deionized water followed by a bath in ethanol. The coverslips are then dried using dry air. A solution containing 484 µl of acetic acid, 56 µl of Bind-Silane (GE Healthcare, New York, USA) completed up to 15 ml with absolute ethanol is prepared. 200 µl of this solution is pipetted to each coverslip and wiped off after few tens of seconds with a dust free wiper. This step is essential to allow stable, covalent bonding of the polyacrylamide hydrogels onto the glass coverslips.

A droplet of 30 µl of the photosensitive solution of polyacrylamide monomers is deposited onto an activated 20 mm coverslip and sandwiched with the grey level mask. UV illumination (Eleco UVP281, Gennevilliers, France, 2W/cm^2^) polymerizes the polyacrylamide. The hydrogel is then gently detached from the grey level mask with tweezers after soaking it into deionized water for 3 min. In this study, the exposure times and chrome thicknesses that we used to control the hydrogel rigidity were respectively 20 s and 29 nm for the millimeter-scaled rigidity pattern (“concentric” design, Fig. 1A), 100 s and 39 nm for the centimeter-scaled rigidity pattern (“double rigidity” design, Fig. 1B), and 20 s and 14 nm for the micrometer-scaled rigidity patterns (“stripes” design, Fig. 1C).

**Figure 1:**
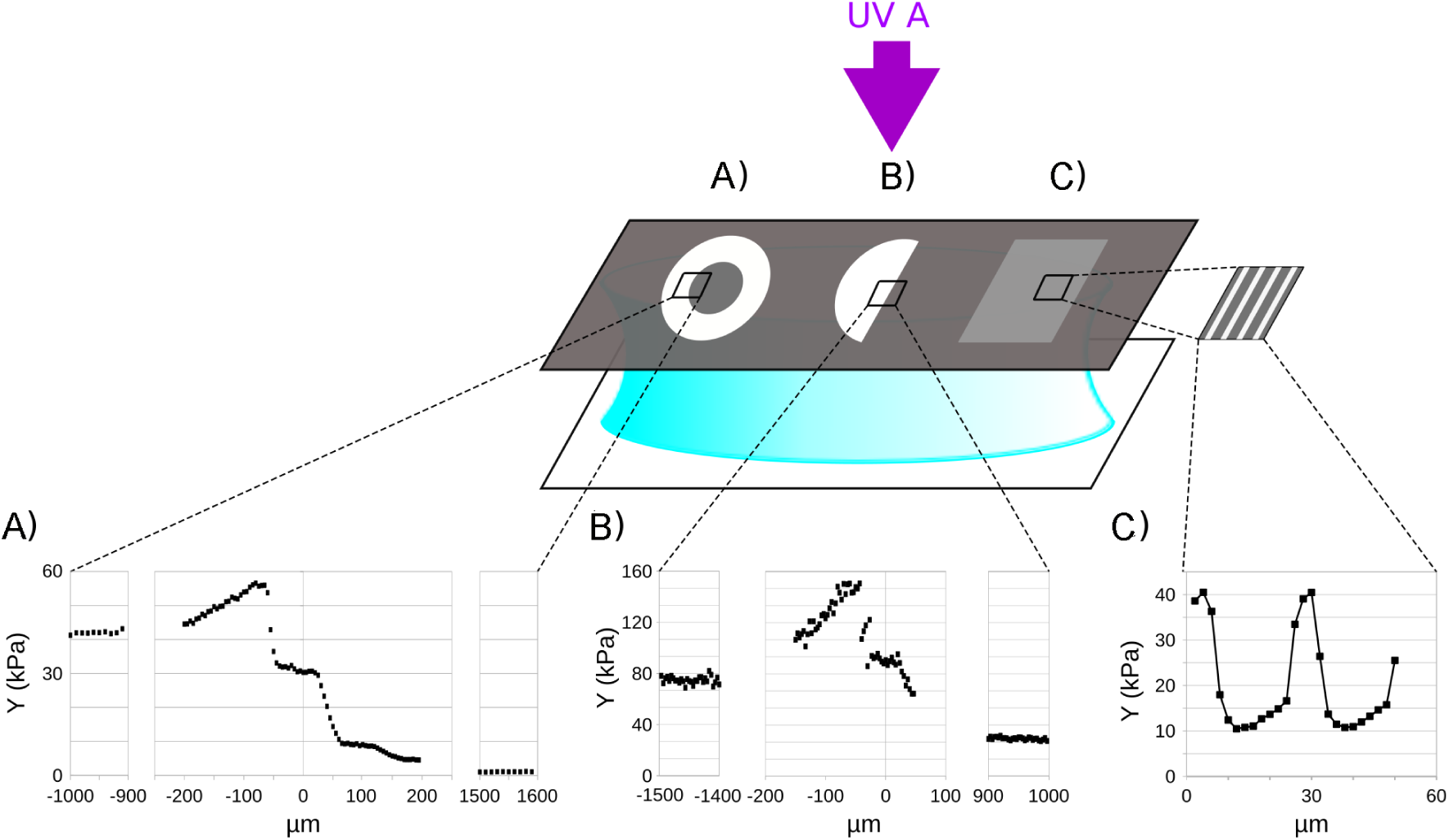
Technology for the fabrication of the culture supports and designs of the rigidity patterns. A) “Concentric” design, millimeter-scaled pattern. Young’s modulus Y of 1.1 ± 0.2 kPa (soft) and 42.0 ± 1.1 kPa (stiff). B) “Double rigidity” design, centimeter-scaled pattern. Y of 29.2 ± 1.1 kPa (soft) and 74.8 ± 2.8 kPa (stiff). C) “Stripes” design, micrometer-scaled pattern. Y (and stripe width) of 12 kPa (20 µm) (soft) and 40 kPa (5 µm) (stiff). Quantification of the substrate stiffness is presented as mean ± standard deviation (SD).

Hydrogel stiffness was characterized by measuring the Young’s modulus using an atomic force microscopy in the force mapping mode (NanoWizard II, JPK Instruments AG, Berlin, Germany). MLCT C tips cantilevers (Bruker, Santa Barbara, USA) with a nominal spring constant of 0.01 N m^−1^ have been employed for their ability to address Young moduli between 0.1 and 50 kPa (Fig. 1).

Hydrogels were functionalized using Sulfo-LC-SDA photosensitive reagent (Sulfo-NHS-LC-Diazirine, Pierce Biotechnology, Waltham, USA), according to a previously reported protocol (45). We employed fibronectin (FN, from human plasma, Roche Applied Science, Switzerland, ref 11080938001) or poly-L-lysine and laminin (PLL, P2636; LN, L2020, Sigma-Aldrich, St. Louis, USA). Hydrogels were first dehydrated until the surface looks dry. Covalent grafting of the FN was performed as follows: FN was first coupled to the heterobifunctional crosslinker sulfo-LC-SDA (Pierce, Waltham, USA) (45). The dehydrated gels were then incubated with the solution of diazirine-coupled FN at concentration 3.5 µg/cm^2^ for 1 h (FN coating). The solution was then drawn off and the hydrogel was immediately exposed to UV illumination (Eleco UVP281, 2 W/cm^2^) for 5 min so that the diazirine function of the sulfo-LC-SDA binds to the surface of the polyacrylamide hydrogel. Covalent grafting of PLL and LN was performed in two steps. 300 µl of a solution of sulfo-LC-SDA at concentration 0.44 mg/ml in water was deposited on the dehydrated hydrogels for 2 h. The solution was then pipetted off and the hydrogels dehydrated again. Sulfo-LC-SDA was covalently bound to the hydrogel by UV illumination for 5 min. 300 µl of a solution of PLL at concentration 1 mg/ml was then incubated on the hydrogel for 1 h. After removal, 300 µl of a solution LN at concentration of 10 µg/ml was then deposited on the hydrogel and let for react for 1 h. The hydrogels were then rinsed 3 times in phosphate-buffered saline (PBS).

The characterization of the surface coating was performed using immunostaining. The following antibodies and respective secondary antibodies were used in the indicated dilutions: polyclonal FN or LN antibodies (Sigma-Aldrich, St. Louis, USA, F3648 and L9393 at 1:400) and Alexa Fluor 488 conjugated donkey anti-rabbit (Thermo Fisher Scientific, Waltham, USA, 1:2000). Surface densities were quantified using confocal microscopy (Leica, Wetzlar, Germany, TCS SP2, objective 40x WI NA 0.6). In order to guarantee comparable conditions, the gain was maintained constant during all acquisitions.

### Primary cell culture

Cortical and hippocampal cells were obtained from embryos of 18 days of gestation of C57 BL/6J mice (Charles River, Wilmington, USA). The experiments in this study were approved by the local ethic committee. At this embryonic stage, neurogenesis is almost completed and astrocytes start to differentiate from multipotent precursors (the relatively lower proportion of oligodendrocytes only appear postnatally). The timing of in vivo cell differentiation continues in vitro, leading to a dominant proportion of astrocytes versus other glial cells in our culture (46). Mixed neuronal-glial cell cultures were conducted as follows: after dissection, cortices (including the hippocampus) were dissociated in DMEM medium supplemented with 10% fetal bovine serum (FBS, Thermo Fisher Scientific, Waltham, USA), 1% L-glutamine, 1% sodium pyruvate and 0.05% peni-streptomycine (Thermo Fisher Scientific, Waltham, USA), a medium that we denote DMEMs (supplemented DMEM). Cells were plated on the coated hydrogels at 150 cells/mm^2^ density. After 3 h, the culture medium was renewed and cells were maintained either in neurobasal medium supplemented with 2% B27, 1% L-glutamine and 0.05% penicillin-streptomycin (Thermo Fisher Scientific, Waltham, USA), referred as NBs medium (supplemented NB) and known to optimize neuronal survival (47), or in in DMEMs medium, which is the medium of reference for astrocyte cultures (48). Pure glial cell cultures were conducted as follows: after dissection and tissue dissociation, approximately 3 · 10^6^ cells were plated per 100 mm Petri dish, previously functionalized with a poly-DL-ornithine coating (0.2 µg/cm^2^ solution incubated over night, P8638, Sigma-Aldrich, St. Louis, USA). This medium improves glial cell proliferation whereas favoring neuronal death, thus leading to a pure glial cell population. DMEMs medium was renewed at 1 and 3 days in vitro (DIV). Cells usually reached confluence at 7 DIV. Cells were then gently detached after 0.05% Trypsin-EDTA (Thermo Fisher Scientific, Waltham, USA) incubation, re-suspended in DMEMs medium and plated (30 cells/mm^2^) on the coated hydrogel substrates. Cells were plated at a low density to limit physical contact in between astrocytes, but with a high enough density to ensure cell survival. Note that this density is much above the number of astrocytes plated in mixed cultures, as the proportion of astrocytes versus neurons for E18 mouse embryos is about 2-3% (49). Consistently, we obtained a similar proportion of glial cells on FN or PLL/LN coated hydrogels, where we quantified cell initial adhesion in serum-free NBs medium (Fig. S2).

### Immunofluorescent stainings of cells

Cells were fixed in 4% paraformaldehyde for 20 minutes and immunostained with standard techniques, after a permeabilization step of 30 min in PBS supplemented with 2% bovine serum albumin (BSA) and 0.25% Triton X-100. The following antibodies and respective secondary antibodies were used in the indicated dilutions: for astrocytes cytoskeleton, Rabbit anti GFAP antibody (kindly supplied by A. Triller, ENS Paris, 1:250) and Alexa 488 conjugated goat anti-rabbit (Thermo Fisher Scientific, Waltham, USA, 1:250), for neurons, anti MAP2 antibody (Thermo Fisher Scientific, Waltham, USA, 1:500) and CY-3 conjugated goat anti-mouse (Thermo Fisher Scientific, Waltham, USA, 1:300), for actin, phalloidin Alexa Fluor 594 (Thermo Fisher Scientific, Waltham, USA, 1:150), for cell-cell adhesions, N-cadherin (kindly supplied by D. Gulino-Debrac, CEA Grenoble, 1:200) and Alexa Fluor 488 conjugated goat anti-mouse (1:300), for DNA, Hoechst (Thermo Fisher Scientific, Waltham, USA, 1:1000).

### Imaging

Fixed cells were analyzed by phase contrast or fluorescence with an inverted Olympus CKX41 or a upright Olympus BX51 microscope at 10x magnification. Time-lapse acquisitions were performed with a inverted microscope (Olympus, IX73). equipped with a heated workplate, a humidifier, a CO2 delivery system and a motorized stage to allow multi-position and multi-condition acquisitions.

### Data analysis and statistical analysis

Cells were observed between 2 and 21 days in vitro (DIV) by maintaining the same samples in culture over the entire time span. Mean cell density values were calculated on a number of fields of view (surface of around 0.5 mm^2^) varying from 4 to 12 and depending on the localization on the sample of the area of interest. This corresponds to an amount of counted cells from a few tens to more than 2000 (in cases of high proliferation). These regions were always chosen in diametrically opposed positions on the circular shape of the gel, in order to get representative descriptions of cell distribution. Neurons and glial cells were quantified after medium change, and thus after dead cell removal. Both cell types were identified by their morphology from phase contrast images. Other cell types, for instance microglia, are distinguished, and therefore discarded, by their shape and fast dynamics (Fig. S3). The initial cell adhesion is calculated as the cell density at the first day of observation (2 DIV for pure cultures) normalized by the seeding cell density. The proliferation rate is calculated as the number of cell divisions *n* divided by a given temporal range Δ*t* = *t*_*i*_ − *t*_0_. The number of cell divisions *n* is obtained by the following equation: *N*(*t*_*i*_)/ *N*(*t*_0_) = 2^*n*^, where N is the average cell density. The mean cell surface is calculated by dividing the total area of fields of view with confluent cells (i.e. with no free-surface) by the number of adherent cells. Statistical analyses and linear regression were carried out with GraphPad Prism software. A Mann-Whitney test (two-tailed, unpaired, with a significance level of 0.05) was performed between two groups of non-normal data.

## RESULTS

In the following, we use the term ‘glial cells’ to talk about the non-neuronal cells that are predominantly mature astrocytes, as a consequence of the stage of development of the embryos and the culture methods that we employed (4, 50).

### The density of the surface coating is independent of rigidity gradients

In order to address primary brain cells sensitivity to rigidity and gradients of rigidity, using grey leveled lithography technique, we designed polyacrylamide hydrogels with centimeter, millimeter and micrometer-scaled apposed regions with distinct pairs of rigidities, e.g. 1.1kPa and 42kPa (or 29kPa and 75kPa) which we thereafter named “soft” and “stiff” areas, respectively (Fig. 1). We could check that the surface density of the adhesion proteins did not show any significant variations with the rigidity even in the regions where the gradient of rigidity is of order of few kPa/µm (Fig. S4). Topography was observed on centimeter and millimeter-scaled rigidity patterns in the place of the gradient of rigidity (Fig. S4A). The transition zone between soft and stiff large areas (millimeter to centimeter-scaled) displayed a smooth topography consisting in a step of a most a 25 µm step over a length of 40 µm (Fig. S4A). The profile of the surface did not evidence sharp angles, which are known to influence cell shape and adhesion (51, 52). Instead, concave curvatures were observed on the soft side of the border between the pairs of rigidities. However, a preferential localization of the cells at these places was not observed, as could result from curvature-induced cell positioning (53). Consequently, cell behavior at the frontier between the stiff and the soft zones were attributed to the presence of rigidity gradients and not to topography. Finally, no topography was observed on micron-scaled patterns of rigidity (Fig. S4B).

### Neuron adhesion depends on the rigidity whereas survival does not

Neurons sensitivity to rigidity was probed on “concentric” millimeter-scaled patterns Young’s moduli of 1.1 ± 0.2 kPa (soft) and 42.0 ± 1.1 kPa (stiff) (Fig. 1A). Neuron adhesion and survival until 17 DIV was analyzed from cells grown in mixed cortical culture in NBs in the absence of serum as described in the Materials and Methods (Fig. 2). Poly-L-Lysine/Laminin (PLL/LN) coating was employed to ensure neuron adhesion as fibronectin (FN) did not allow significant neuron attachment and spreading (Fig. S5). It was observed that, although the amount of adherent neurons was slightly but significantly larger on the soft substrates than on the stiff ones (Fig. 2C), neuron density on both rigidities appeared statistically constant over the entire time span, thus showing that neuron survival is independent of the rigidity of the substrate. The slight variations in the number of neurons over time was attributed to the difficulty to count them in bright field images, especially after two weeks of culture when neurons are organized in mature networks and where individual somas are difficult to identify.

**Figure 2:**
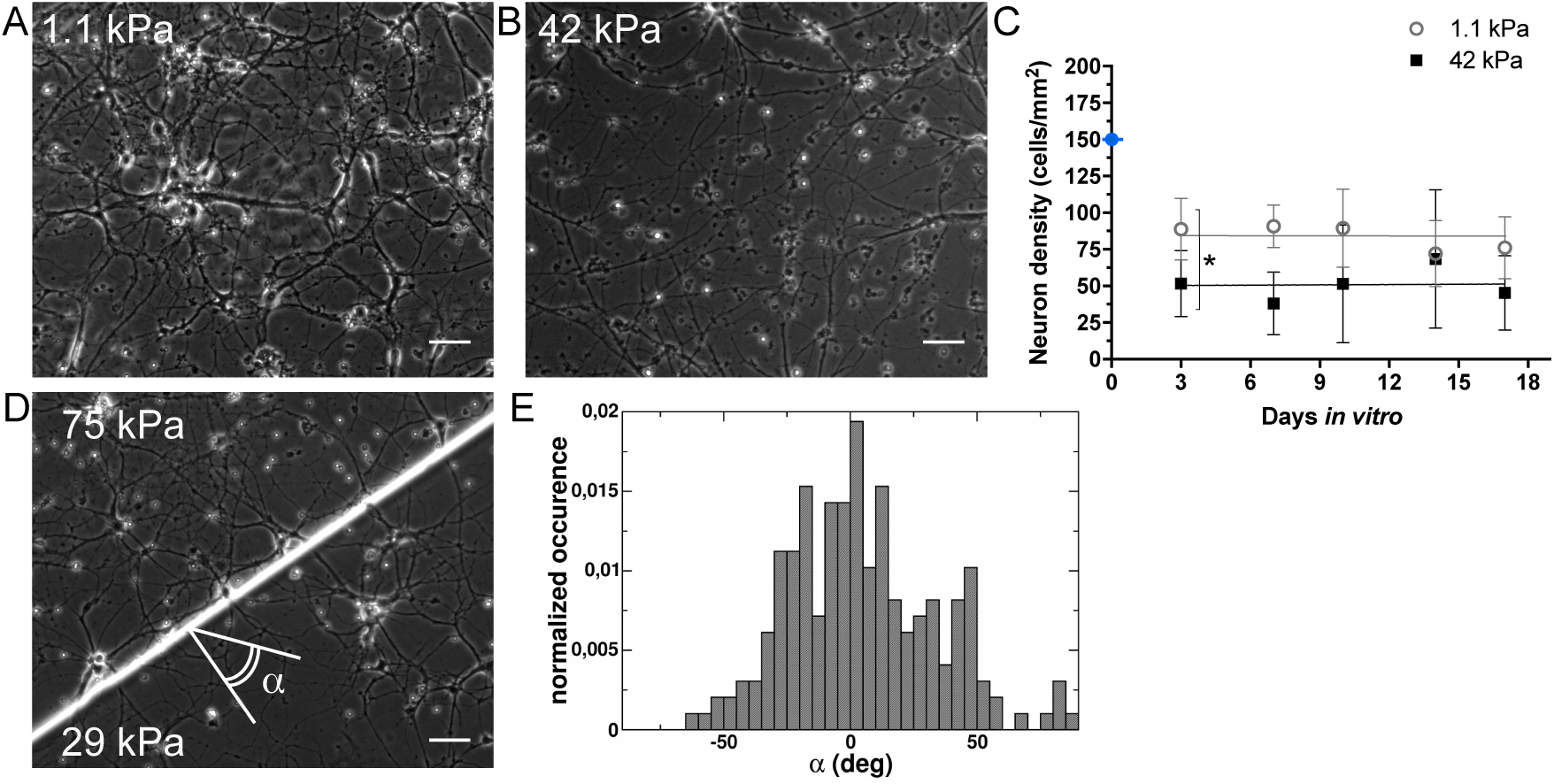
Neuronal cell growth in mixed cultures on laminin/Poly-L-lysine coated substrates in NBs culture medium. A, B - Phase contrast imaging of neuronal cells on a soft (A, 1.1 kPa) and high stiffness (B, 42 kPa) on “concentric” pattern of rigidity at 14 days in vitro (DIV). Scale bars: 50 µm. C - Neuron density and its evolution over time (DIV) on stiff (black squares) and soft (white circles) regions. The blue dot indicates the initial cell concentration (including both glial and neuronal cells) seeded on the substrate. Quantification of the cell density is presented as mean ± standard deviation (SD). * denotes that the two means are significantly different, p (stiff vs. soft at 3 DIV) = 0.0380. Slopes not statistically different from zero: p (soft) = 0.1502; p (stiff) = 0.6781. The grey and black lines in the graphes represent the y-mean value of the linear fit. D - Phase contrast imaging of neurons at the frontier of between the low and the high stiffness regions on the “double rigidity” design. E - Neurite orientation at the frontier between the soft and the stiff regions, characterized by the angle *α* perpendicular to the frontier as shown in D. The histogram for *α* is centered at 0 deg.

### Neurites grow preferentially parallel to kPa/µm gradients of rigidity

The orientation of the neurites at the border between the stiff and the soft regions was quantified by analyzing the phase contrast images of mixed cortical cultures grown on a “double rigidity” pattern (Fig. 1B), with Young’s moduli of 29.2 ± 1.1 kPa and 74.8 ± 2.8 kPa (Fig. 2D). The orientation of 196 neurites from 3 experiments was analyzed. We observed that the neurites predominantly cross the rigidity border perpendicular to it, thus aligning with the gradient of rigidity (Fig. 2D,E).

### Glial cell adhesion and proliferation in pure cultures are favored on high rigidity in the presence of fibronectin coating

As glial cells do not adhere nor proliferate properly in culture conditions that are favorable to neurons (PLL/LN coating and NBs culture medium, Fig. S2B), glial cells sensitivity to rigidity was first evaluated in pure culture conditions. Glial cells were seeded on fibronectin (FN) coated- “concentric” rigidity patterns (Fig 1A) characterized by Young’s moduli of 1.1 ± 0.2 kPa (soft) and 42.0 ± 1.1 kPa (stiff), and cultured in DMEMs. The cell density was measured from 2 to 17 days in vitro (DIV) (Fig. 3). We observed an almost absence of cells on the soft region during the entire time span (Fig. 3B). Residual cells kept round and did not spread. Conversely, on the stiff regions, the initial cell density of seeding (around 30 cells/mm^2^) was conserved after plating, showing that the combination of a 42 kPa stiff substrate and FN coating supports the adhesion and survival of astrocytes. The proliferation rate was close to constant between 2 and 8 DIV, with an averaged value of 0.27 (Fig. 3B). Furthermore, the glial cells were widely spread on the stiff regions (mean cell surface: 14.5 ± 3.4 · 10^3^ µm^2^). Confluence was reached between 8 and 17 DIV, at a plateau of cell density of average of 60 ± 22 cells/mm^2^ (Fig. 3B).

**Figure 3:**
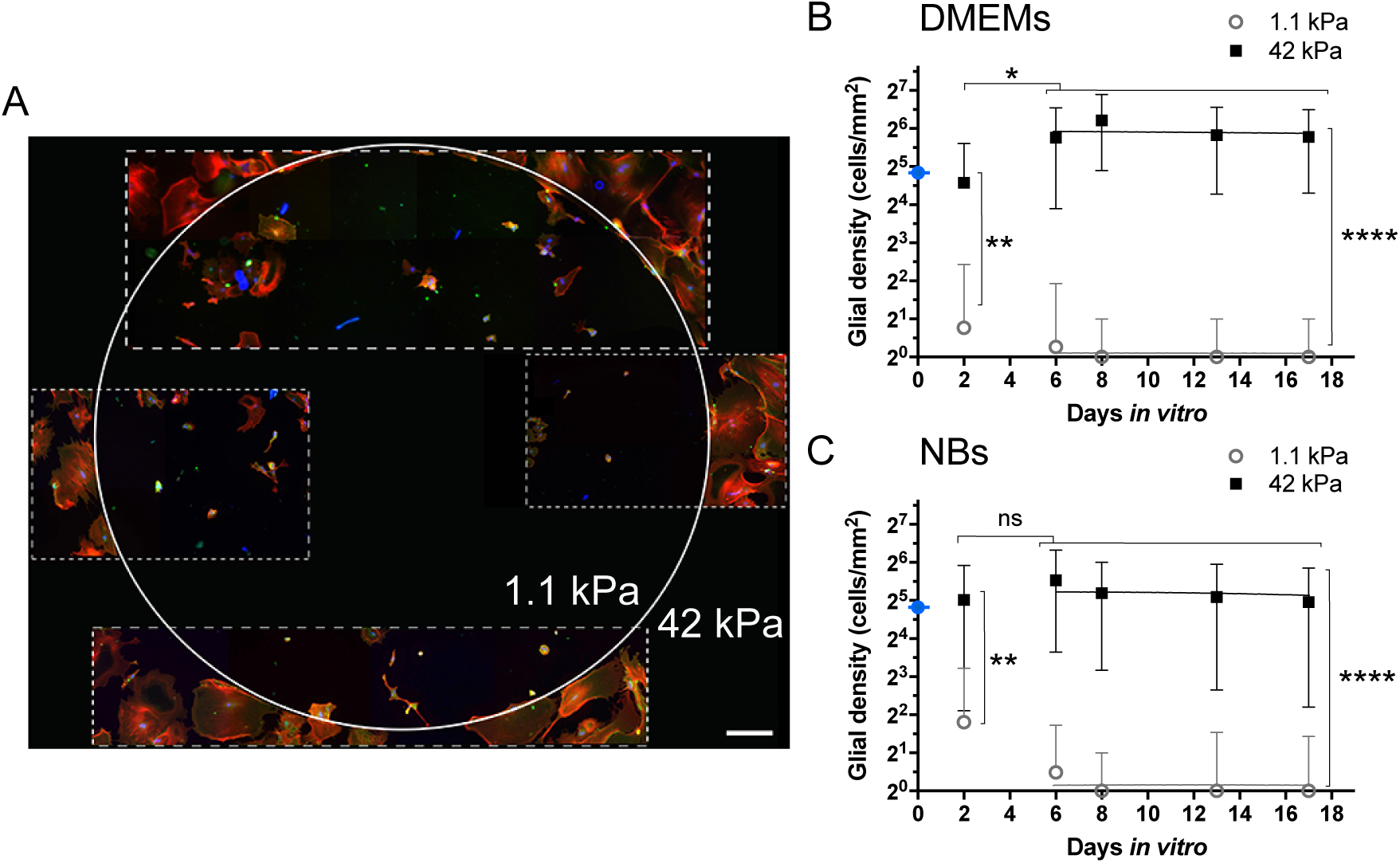
Glial cell initial adhesion and proliferation in pure cultures on fibronectin coated substrates. A - Immunofluorescence images of a glial cell culture in DMEMs medium: the frontier (solid white line) delineates the soft, inner region and the stiff, outside area. The dashed white lines mark out the immunofluorescence images in the rigidity pattern. Red: phalloidin, actin. Green: N-Cadherin, cell-cell adhesions. Blue: Hoechst, nuclei. Cells are fixed at 12 DIV. Scale bar: 300 µm. B - Evolution of the cell density in DMEMs medium over time (days in vitro, DIV) on stiff (black squares) and soft (white circles) regions. ** denotes that the two means are significantly different, p (stiff vs. soft) = 0.0050 (2 DIV); *, p = 0.0187 (stiff, 2 DIV vs. between 6 and 17 DIV); ****, p < 0.0001 (stiff vs. soft between 6 and 17 DIV). Slopes not statistically different from zero: p = 0.6463 (stiff) and 0.3288 (soft). C - Evolution of the cell density in NBs medium over time, with identical conventions as in B. **, p (stiff vs. soft at 2 DIV) = 0.0094; ns, p = 0.6207 (stiff, 2 DIV vs. between 6 and 17 DIV); ****, p < 0.001 (stiff vs. soft between 6 and 17 DIV). Slopes not statistically different from zero: p = 0.1164 (stiff) and 0.3288 (soft). The grey and black lines in the graphes represent the y-mean value of the linear fit. The blue dots indicate the initial cell concentration seeded on the substrate. Quantification of the cell density is presented as mean ± standard deviation (SD).

### Glial cell rate of proliferation is dependent of the culture medium and of the presence of serum

In order to study the impact of the culture medium and of the serum, glial cells from pure culture conditions were grown as above on “concentric” 1.1-42 kPa rigidity patterns coated with FN coating but using NBs as culture medium instead of DMEMs (the referenced medium for this kind of culture, see (48)). In order to adhere, cells were first grown for 24 h in DMEMs medium. DMEMs was then replaced by NBs for the following days. As above, a weak adhesion and a poor survival were observed on the soft region. Contrarily to Fig. 3B, Fig. 3C shows that the proliferation rate on the stiff region is almost null, the slope of the cell density over the time between 2 and 17 DIV being non-significantly different from zero (p = 0.5366). Thus, the culture medium, including the presence of serum, is critical to obtain a significant rate of proliferation on the stiff substrate.

### Changing fibronectin coating to laminin/Poly-L-Lysine reverses glial cell response to rigidity

Surface coating was changed to PLL/LN, a surface coating that is less favorable to glial growth (48). Glial cells were grown on identical pattern of rigidity as above using pure culture conditions in DMEMs. Strikingly, the dependence on the rigidity of the glial cell adhesion and proliferation was reversed using PLL/LN instead of FN (Fig. 4A, 6A, B). Indeed, only rare cell adhesion and almost no proliferation were observed on the stiff regions (Fig. 4A). Conversely, cells adhered on the soft regions and a weak but significant cell proliferation was measured, associated to widely spread morphologies similar to those observed on FN coating (mean cell surface: 14,3 ± 4.3 · 10^3^ µm^2^). Cell proliferation rate on the soft regions reached a value of 0.4 between 2 and 6 DIV. After 8 DIV, cell density reached a plateau associated to a cell confluence of approximately 60% (Fig. 4A). We investigated whether this observation was conserved in a different culture condition. Mixed cortical cultures were thus grown on identical culture supports as above in DMEMs. In this condition characterized by a very small initial proportion of glial cells compared to neurons (about 2-3% (49)), we also observed that the PLL/LN adhesive coating promotes a larger proliferation rate of glial cells on soft substrates than on stiff ones (Fig. 4B, 6B). However, contrarily to the case of pure culture, a non-zero proliferation rate is observed on stiff regions.

**Figure 4:**
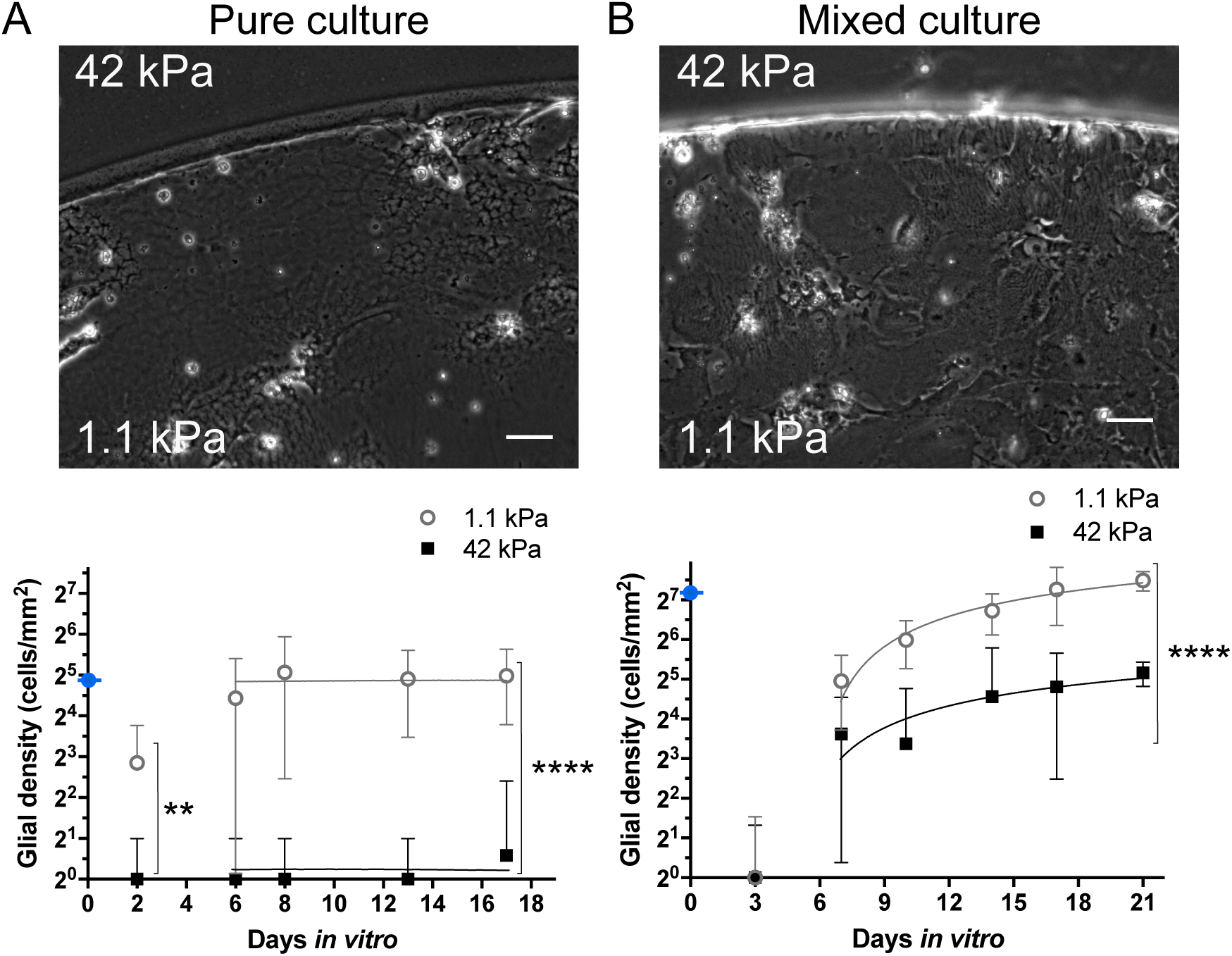
Glial cell initial adhesion and proliferation on poly-L-lysine/laminin coated substrates in DMEMs culture medium. A - Glial cells from pure culture at the stiff (42 kPa) - soft (1.1 kPa) frontier on a “concentric” pattern of rigidity at 17 DIV, and the evolution of the cell density over time (days in vitro, DIV) on stiff (black squares) and soft (white circles) regions. ** denotes that the two means are significantly different, p (stiff vs. soft) = 0.0016 (2 DIV); ****, p < 0.0001 (between 6 and 17 DIV). Slopes not statistically different from zero: p = 0.1946 (stiff) and 0.4699 (soft). The grey and black lines in the graphes represent the y-mean value of the linear fit. B - Idem for glial cells from mixed culture (phase contrast image at 17 DIV). ****, p < 0.0001 (stiff vs. soft between 6 and 17 DIV). Black (stiff) and grey (soft) lines represent semilog fits. The blue dots indicate the initial cell concentration (including both glial and neuronal cells for the mixed culture) seeded on the substrate. Quantification of the cell density is presented as mean ± standard deviation (SD). Scale bars: 50 µm.

### Glial cell adhesion and proliferation in mixed cultures show an enhanced response to rigidity compared to pure culture, associated to limited spreading

The proliferation rates of glial cells from pure and mixed culture conditions were both enhanced on the soft region on PLL/LN coating (Fig.4). However, quantitative differences in cell adhesion or proliferation rates were observed (Fig. 6). We therefore investigated whether glial cell adhesion and proliferation would be enhanced by the presence of neural cells also on FN coating, even if neurons do not survive long in these medium and coating conditions. Cells were grown on the “concentric” pattern (Fig. 1A) with FN coating and DMEMs culture medium, so to allow glial cell proliferation and progressive neuronal death. As in pure glial cultures, glial cells in mixed cultures adhered preferentially on stiff regions (Fig. 5A), with cell initial density at 3 DIV being larger by a factor of more than twenty on the stiff than on the soft regions (Fig. 5B). We compared the rates of glial cell proliferation depending on the culture conditions (Fig. 6B). We first observed that mixed culture conditions are more favorable to glial cell adhesion: while almost no cells from pure culture conditions adhere in the presence of unfavorable rigidity (soft on FN, stiff on PLL/LN), a small number of adherent glial cells from mixed culture conditions is sufficient to reach a significant proliferation rate in all conditions (Fig. 6B). Indeed, the proliferation was enhanced for all coatings and rigidity in mixed culture conditions compared to pure cultures (Fig. 6B,C). This led to very dense layers of cells on the stiff regions coated with FN, with their mean surface going down to 3.3 ± 1.8 · 10^3^ µm^2^, more than 4 fold smaller than in pure culture conditions. However, this large rate of proliferation and cellular compaction led to the detachment of entire portions of the cell monolayer, causing the impossibility to count cells later than 10 DIV (Fig. 6C). We conclude that the initial presence of neurons therefore enabled cell adhesion in all conditions and favored cell proliferation. This observation could however not be extended for cells cultured in NBs. As in pure glial culture, the use of NBs instead of DMEMs did not allow to maintain a detectable proliferation, independent of the stiffness and the nature of the adhesive coating (Fig. S2).

**Figure 5:**
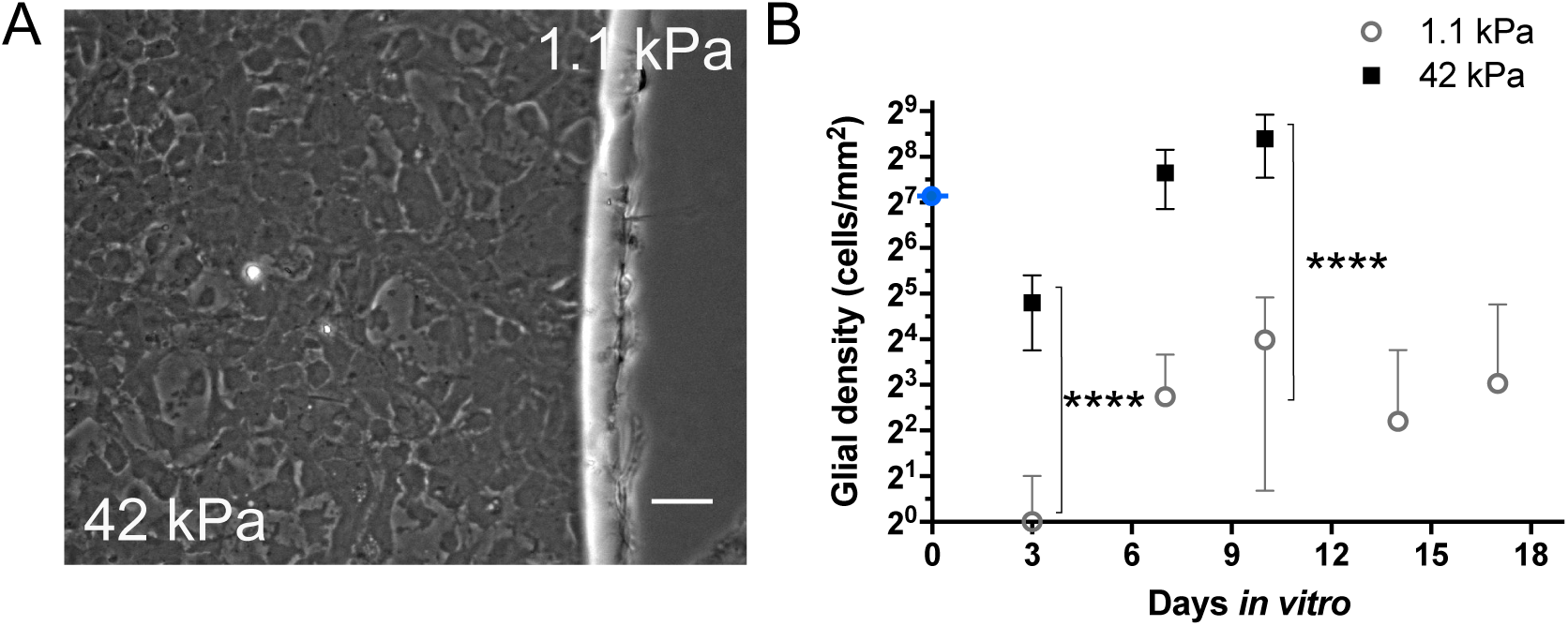
Glial cell initial adhesion and proliferation in mixed cultures in DMEMs medium on fibronectin coating. A - Phase contrast imaging of glial cells at the border of stiff (42 kPa) and soft regions (1.1 kPa) at 21 DIV of a “concentric” pattern of rigidity. Scale bar: 50 µm. B - Evolution of the cell density over time (DIV) on stiff (black squares) and soft (white circles) regions. **** denotes that the two means are significantly different, p (stiff vs. soft at 3 and at 10 DIV) < 0.0001. The blue dot indicates the initial cell concentration (including both glial and neuronal cells) seeded on the substrate. Quantification of the cell density is presented as mean ± standard deviation (SD).

**Figure 6:**
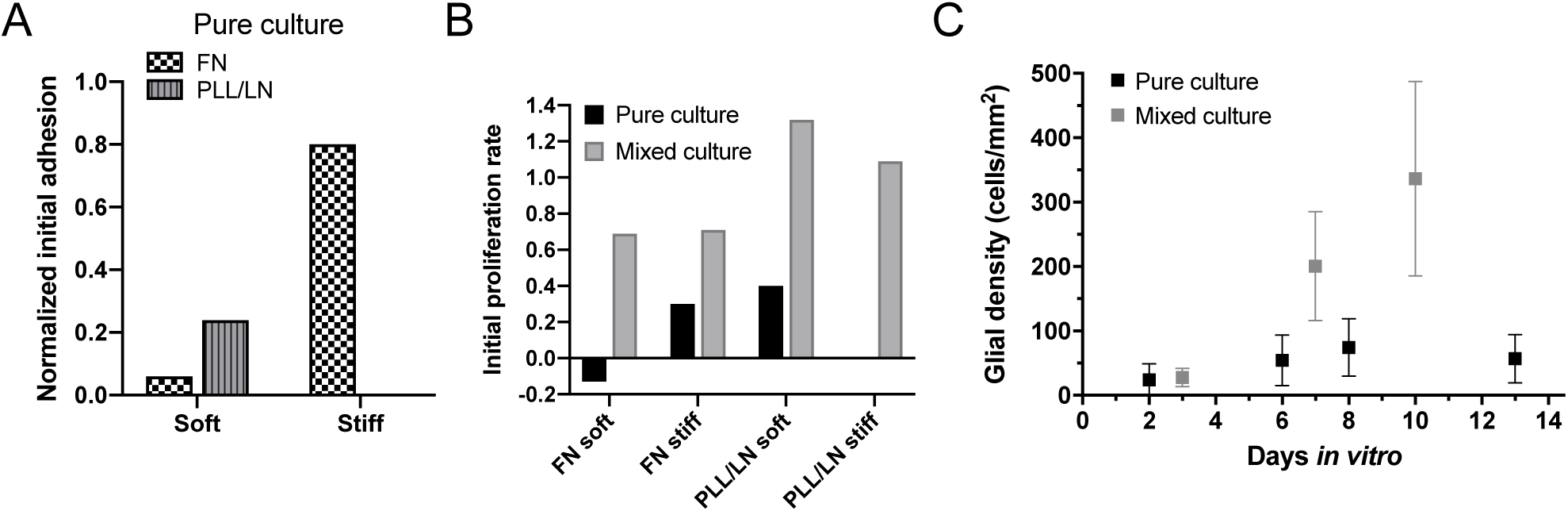
Rigidity sensitivity in DMEMs of glial cells from mixed cultures is enhanced compared to pure cultures. A - Comparison of the normalized initial adhesion of glial cells from pure cultures on the different coatings (“FN”: fibronectin, “PLL/LN”: poly-L-lysine/laminin) and rigidities (“Soft”: 1.1 kPa, “Stiff”: 42 kPa). B - Normalized initial proliferation rates show that glial cells proliferate faster in mixed culture conditions. C - Comparison of the evolution of the cell density in pure and mixed cultures on the stiff regions coated with FN.

### Glial cells adapt their shape to subcellular rigidity gradients

Observation of glial cell shape on centimeter or millimeter-scaled patterns of rigidity showed that glial cells often align at least part of their body with the frontier between the stiff and the soft regions (Fig. 3, 4, 7, Movie S1). We thus wondered how micrometer-scaled patterns of rigidity would impact their adhesion and shape, whether they could get confined by gradients of rigidity. For this purpose, glial cells from pure culture conditions were grown on a parallel array with alternated soft and stiff areas at a scale similar to the cellular sizes (Fig. 1C). More precisely, borders between the two rigidities were separated by distances from 5 to 75 µm. A uniform FN coating and DMEMs medium were used for this study, to optimize cell adhesion and to culture cells in proliferating conditions. We observed that single glial cells get deformed by the rigidity patterns (Fig. 7B, pattern of 5-20 µm stripes, i.e. when stiff stripes are narrower than the soft ones). They preferentially extended lamelippodia on the stiff stripes. But we did not observe cell confinement on the patterns as for other cell types (Fig. S6), even when enlarging the distance between the stiff stripes (Fig. S7).

**Figure 7:**
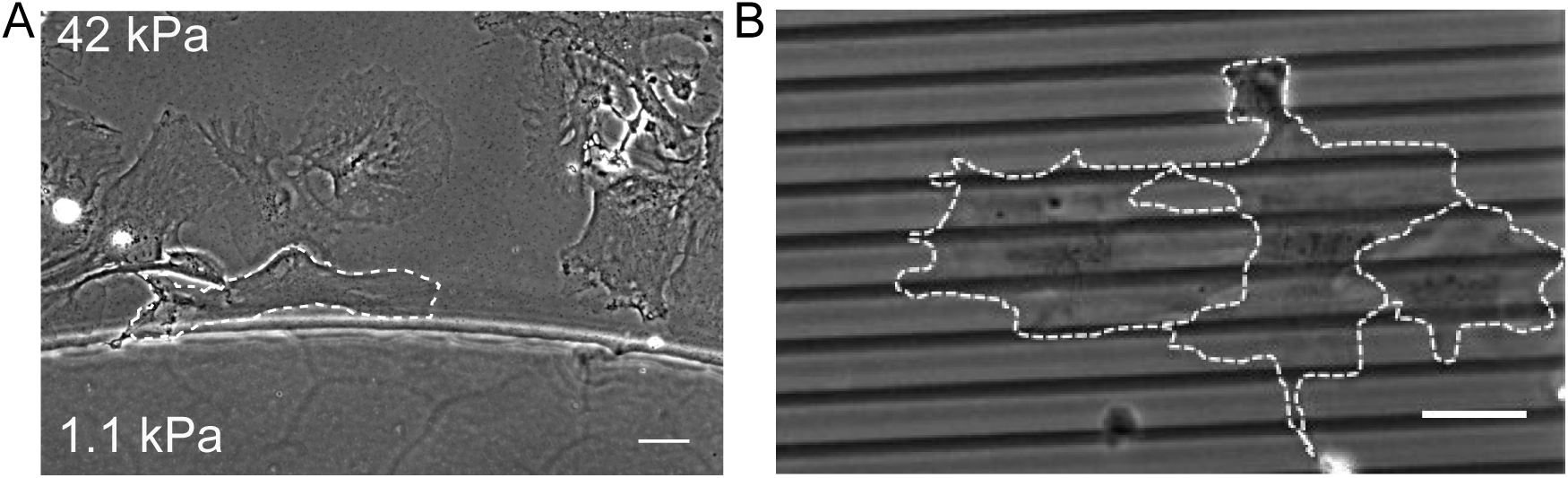
Glial cells from pure cultures in the presence of gradients of rigidity in DMEMs medium on fibronectin coating. A - Glial cells grown on “concentric” rigidity pattern align perpendicular to the gradient of rigidity at the border between the stiff and soft regions (17 DIV). B - Glial cells are deformed by subcellular rigidity patterns (stiff stripes of 5 µm, 40 kPa, alternating with soft stripes of 20 µm, 12 kPa, 2 DIV). Scale bars: 50 µm.

## DISCUSSION

Using functionalized multi-rigidities polyacrylamide substrates, we have investigated the combined response of primary brain cells to the stiffness and chemical coating of their extracellular matrix. This technique has proved to be a unique tool to probe cellular responses to a panel of rigidities under identical culture conditions in terms of culture medium, adhesive molecules and soluble cues secreted by the cells. Two culture media have been tested, one more favorable for glial cells (serum-rich DMEMs), the second one more adapted for hippocampal neurons survival (NBs, serum-free medium). Two coatings have been tested, based on two components of the ECM in the brain: fibronectin (FN) and laminin (LN, associated to poly-L-lysine, PLL) (54). In this study, two types of initial cell preparations have also been investigated (purified glial cell suspension or dissociated cortical tissue), to test the influence of neurons on glial cells mechanosensitivity.

We could first confirm that embryonic neurons in mixed cultures adhere better on softer supports (21). We then observed that their survival is not sensitive to the rigidity. This observation differs from Ref. (21) where neurons on the stiff polyacrylamide hydrogel (27 kPa) were mostly observed over glial cells and not directly on the stiff hydrogel itself. Here glial cells sparsely survived either on the soft or the stiff regions in these culture conditions (NBs medium, absence of serum, PLL/LN coating) (Fig. S2A) while neuron density kept significant on both regions (Fig. 2C). Differences of more than an order of magnitude of the surface density of the coating may be at the origin of this distinct behavior, the viability of neurons being dependent of laminin density (55). Nevertheless, following Ref. (21) and Ref. (30), we also observed a smaller density of neurites on the stiff than on the soft regions (Fig. 2A, B).

We then analyzed the orientation of neurite growth on gradients of rigidity. We clearly observed that neurites align with the gradient of rigidity (Fig. 2D, E). Ref. (56), (57), (43) and (27) reported that neurite length depends on the rigidity of the extracellular matrix and that this response can be further modulated by the coating (27). Ref. (56) moreover showed that neurite growth is faster on a stiff than a soft support, and straighter. This would suggest that neurites could grow upstream or downstream the rigidity profile, thus parallel to the gradient, the growth being speed up and oriented by the stiffer region. Consistent with this analysis, we observed that neurites oriented parallel to the gradient of rigidity. Accordingly, Ref. (43) also reported neurite growth parallel to the gradient of rigidity, from stiff to soft, in 3D collagen gels. Ref. (56) nevertheless reported that axons locally grow perpendicular to the gradient of rigidity, but parallel to it (from stiff to soft) on larger distances (Fig. 6 in (56)). The fact that different scales could cause antagonist responses to rigidity gradients is far from obvious and is not supported by the identification of scale-dependent signaling pathways yet. We therefore suspect that this observation might be related to other physical cues such as a competition between a persistent movement favored on stiff regions and a reorientation caused by the gradient of rigidity. Contrary to Ref. (56) and Ref. (27), we could not confirm that neurite growth is identical on FN and PLL/LN as neurons cultured on FN coating grew on top of glial cells and arranged as 3D aggregates (Fig. S5).

Concerning glial cells, higher proliferation rates were systematically observed in mixed cultures compared to pure ones. This behavior may be induced by factors secreted by neurons, which have been shown to enhance cortical gliogenesis in early stages (58). Another hypothesis is that their progressive death in DMEMs medium may act as a trigger of glial cell proliferation, as suggested by (59).

Our main result is that the differential response to rigidity of glial cells depends on the surface coating. While we observed that glial cell adhesion and proliferation on FN is favored on the stiff regions as already reported (6, 21, 31) (Fig. 3), we observed an opposite behavior using PLL/LN coating (Fig. 4). In this condition, larger adhesion and proliferation occurs on soft regions. This observation was shown to be independent of the cell preparation, glial cells being either purified from neurons before seeding or seeded as cortical, mixed cultures (Fig. 4 and Fig. 5). Contrarily to observations reported on poly-D-lysine coating (50), glial cell purification that was conducted in Petri dishes for a week to promote their growth and to suppress neurons was only shown to impact their capability to proliferate and not their intrinsic adaptation to soft or stiff environments (Fig. 6). We therefore suggest that this behavior may originate from the integrins that are involved in cell adhesion. Integrins subtypes have been shown to differently contribute to rigidity probing (60). For instance traction forces transmitted by *α*_6_ *β*_1_ integrins which are involved in cell adhesion on LN (61) were shown to be of smaller amplitude than those transmitted by the *α*_5_ *β*_1_ integrins on FN (62). LN was also shown to involve a different signaling pathway than FN for the mechanosensitivity of neurons (24). Despite these clues, coating-induced reversed mechanosensing had not been reported yet. Contrarily to glial cells, neurons adhere and grow better on a soft matrix (21) (Fig. 2), and changing the nature of the coating from PLL/LN to FN did not change their preference for soft substrates (Fig. S5). The dependence on the adhesive cues of glial cell mechanosensing may however not be specific to this cell type. A recent study of Stanton et al. (63) pointed that the influence of rigidity in stem cell differentiation depends on the nature and the surface density of the chemical coating. In particular, these authors evidenced that LN coating induces the sequestration of the YAP/TAZ transcriptional factor in the cytoplasm of human mesenchymal stem cells on 3 kPa matrix, contrarily to FN, collagen I or collagen IV on which YAP/TAZ can be translocated to the nucleus at specific surface densities. On stiff substrates (38 kPa), YAP/TAZ translocation was only reported at a specific surface density of LN coating, while the other coatings would induce YAP/TAZ translocation on a large range of surface densities. Taken together with our findings (Fig. 4), these observations could indeed point that the absence of YAP/TAZ translocation on stiff matrices on most laminin coatings comes from the fact that combination of a stiff environment and laminin coating is not favorable to cell survival.

We showed that glial cells sense rigidity by analyzing the dependence of their adhesion and proliferation with rigidity. Glial cells from E18 mice embryos are not migratory cells, thus durotaxis cannot be probed. However, immature astrocytes are motile cells and take part to the formation of the central nervous system or to wound repair (64). They for instance migrate along axon tracts in the white matter (65). Axons were shown to be stiffer than the extracellular matrix in brainstem (66). This result can probably be generalized to other brain tissues as axons are framed by microtubules that are stiff polymers. Then we wondered whether glial cells could be confined on stiff tracks on FN coating, reminiscent from their migration as immature glial cells on stiff tracks. Surprisingly, we could not observe such confinement on stiff tracks (Fig. 7). Stiff tracks would only impact cell shape by orienting lamelipodia. Varying the width of the stiff stripes did not allow to confine the cells (Fig. S7). This result is surprising as all the other cell types we have manipulated indeed got confined when the width of the stripes would be of the order of the size of the nucleus (Fig. S6). Further investigations of glial cell mechanosensitivity at subcellular scale is therefore required.

## CONCLUSION

Using multi-rigidity polyacrylamide substrates with rigidity-independent adhesive coatings, we evidenced for the first time that rigidity-favorable culture conditions may be reversed by changing the nature of the adhesive coating. Thus, primary glial cells from mouse embryo demonstrated a better adhesion and proliferation on stiff substrates when coated with fibronectin, while soft substrates were preferred when the coating consisted in poly-L-lysine/laminin. This observation was not dependent on cell preparation. We therefore suggested that this reversal in cell response to rigidity is due to the specificity of the integrin subtypes that are involved in glial cell adhesion, which depend on the adhesive coating. In addition, glial cells showed an unexpected response to subcellular gradients of rigidity. Contrary to other cell types, glial cells could not get confined on rigidity patterns on the panel of geometries and rigidities we probed. These unique and unexpected behaviors, reversal of rigidity preference with adhesive cues and absence of rigidity-induced confinement, suggest that glial cells may own a very specific machinery for rigidity sensing, that differs from most other cell types. Regarding the involvement of glial cells in brain tissue repair and in cancer, deciphering the pathways that lead to their specific response to mechanical cues is crucial for a comprehensive understanding of their behavior in both physiological and pathological situations.

## AUTHOR CONTRIBUTIONS

C. T., C. V. and A. N. designed the research. C. T. and C. M. carried out the experiments. D. F. engineered the masks. C. T. and A. N. analyzed the data. C. T. and A. N. wrote the article.

## ACKNOWLEDGMENTS

C.T. and C.V. are grateful to A.Triller and S. Colasse for graciously receiving us at the IBENS (Paris) and transmitting their expertise in primary brain cell culture. C. T. was funded by French Ministry of Research. C. M. acknowledges the support by CEA TechnoSanté program. C. T, C. V. and A. N. are indebted to D. Gulino-Debrac for providing animal facilities. This work was supported by ANR (ANR-12-JSVE5-0008). C.M. is founder and CEO of the spin-off Cell&Soft which sells pre-coated soft hydrogels for cell culture with uniform or textured rigidity. A.N. is co-founder and shareholder of Cell&Soft. A. N. is scientific adviser of Cell&Soft.

